# Hypertrophied human adipocyte spheroids as *in vitro* model of weight gain and adipose tissue dysfunction

**DOI:** 10.1101/2021.01.06.425629

**Authors:** Anna Ioannidou, Shemim Alatar, Matilda Åhlander, Amanda Hornell, Rachel M. Fisher, Carolina E Hagberg

## Abstract

The rise in obesity prevalence has created an urgent need for new and improved methods to study human adipocytes and the pathogenic effects of weight gain *in vitro*. Despite numerous studies showing the advantages of culturing adipocyte progenitors as 3D structures, the majority continue using traditional 2D cultures which result in small, multilocular adipocytes with poor representability. We hypothesized that providing differentiating pre-adipocytes with a vascular growth niche would mimic *in vivo* adipogenesis and improve the differentiation process. Here we present a simple, easily applicable culture protocol that allows for the differentiation and culturing of human adipocytes with a more unilocular morphology and larger lipid droplets than previous protocols. We moreover offer a protocol for inducing adipocyte enlargement *in vitro*, resulting in larger lipid droplets and development of several key features of adipocyte dysfunction, including altered adipokine secretions and impaired lipolysis. Taken together, our hypertrophied human adipocyte spheroids offer an improved culture system for studying the cellular and molecular mechanisms causing metabolic dysfunction and inflammation during weight gain.

## Introduction

The prevalence of obesity is rapidly increasing worldwide, with great socio-economic consequences. White adipose tissue is the major lipid storage site. During periods of excess caloric intake, the adipose tissue expands both through hyperplasia, the differentiation of new adipocytes from progenitors, and through hypertrophy, enlargement of existing adipocytes, with the latter dominating during weight gain in adult humans^1,2^. However, with obesity, the expanded adipose tissue also becomes dysfunctional, characterized by a persistent low-grade inflammation, altered adipokine secretion and increased lipid release (lipolysis) in the basal, unstimulated state^3–6^. Due to the hypertrophic nature of the weight gain, increased adipocyte size is another characteristic of adipose tissue dysfunction and disease susceptibility, and associated with metabolic disease even in the lean state^2,7^. Together, these changes greatly impact systemic metabolism by promoting inflammation, dyslipidemia and insulin resistance. It is therefore vital to understand the underlying molecular mechanisms of obesity and adipose tissue dysfunction, which will enable the development of improved treatments against obesity-induced metabolic disease^8^.

Mechanistic research into adipocyte dysfunction has been hampered by the adipocytes’ large size and high lipid content, making them notoriously challenging to work with^9^. Human mature white adipocytes within the subcutaneous fat depot, comprising over 80% of body fat, are unilocular (containing a single lipid droplet) with the nucleus squashed to the side of the cells, typically measuring 63.4 μm (range 59.0–72.1 μm^10^) in diameter in lean individuals which can rise up to 200 μm during obesity. The adipocytes are moreover closely juxtaposed to each other within the tissue and interspersed by a network of micro vasculature and tissue-resident immune cells. Due to their lipid content, mature white adipocytes cannot be frozen, and therefore any culture models such as the recently developed MAAC model require a constant supply of fresh surgical biopsies, not always available^11^. The research community has therefore reverted to using committed adipocyte progenitors, so-called pre-adipocytes, which easily can be differentiated in culture *in vitro*^12^. Differentiation yields adipocytes which readily accumulate lipid droplets and express all the characteristics of human mature cells, but which are unable to become unilocular using the current 2D culture systems, and do not share an *in vivo*-like tissue architecture with juxtaposed cells. The lack of lipid droplet unilocularity is important, as small lipid droplets have a very different membrane-to-volume ratio compared to large ones, and hence will differ in their lipid mobilization dynamics^13^. Moreover, these differentiated cells have been used mostly to study the effects of genes or agents on adipogenesis (adipocyte differentiation), while they have been used much less for the study of mature adipocytes, which is their predominant cellular state *in vivo*.

The cellular niche might be a factor explaining, at least in part, why *in vitro* differentiation has remained poor. Seminal studies in mouse showed that differentiating adipose tissue develops along the vascular tree, using the microvasculature as growth niche^14,15^. Moreover, cell-cell and cell-matrix contacts have also been shown to be essential for differentiation^16^. Reasoning that the lack of adipocyte maturity in current 2D culture models might be due to such key differentiating signals being absent during the differentiation process, we hypothesized that by providing pre-adipocytes with both cell-cell contacts and a vascular network, we would be able to construct an improved human adipocyte culture model. Here we present an optimized spheroid model, culturing and differentiating human stromal vascular fraction (SVF) cells into mature adipocytes imbedded in Matrigel, which yields larger lipid droplets and a higher proportion of unilocular adipocytes than previous methods. We moreover present a simple protocol to ‘fatten’ the spheroids in culture, yielding spheroids with enlarged lipid droplets, altered adipokine signalling and higher basal lipolysis, thus mimicking the phenotypic characteristics of weight-gain and adipocyte dysfunction *in vitro*.

## Methods and Material

### Cell culture

Primary human subcutaneous SVF cells (including preadipocytes, endothelial cells and immune cells) from healthy donors (Lonza, US) were seeded in a 96-well ultra-low attachment (ULA) plate with a round bottom (Corning, Costar #CLS7007) at 10,000 cells per well. On day 6 the formed spheroids were embedded in 40 μL growth factor-reduced (GFR-)Matrigel (Corning, #11553620) and cultured for a total of 40 days. From day 0-10 the cells were seeded and cultured in 200 μL/well Endothelial growth medium 2 (EGM-2, CC-3162, Lonza) to preserve endothelial cell integrity^17^. On day 10, half of the cell culture media was replaced by 2x Preadipocyte Growth Medium (PGM-2, PT-8002, Lonza) and the removed media used for assays. PGM-2 only was used thereafter, adding 50 μL PGM-2 medium on d14, d24 and d34, and removing 150 μL of the media on d20, d30 and d30 to be used for assays, replacing it with fresh PGM-2. To mimic weight gain, spheroid media was supplemented with Intralipid (L0288, Sigma, Aldrich) diluted 1:250 to a final concentration of 800 μg Intralipid per mL media. Analysis and harvesting of spheroids occurred on days 6, 10, 20, 30 and 40. On these days, pictures of the spheroids were taken using a light microscope, media was collected and added back to the wells as described above. In addition, a subfraction of the spheroids themselves was harvested at each time point for analysis for DNA content.

### Microscopic analysis, immunofluorescence and confocal microscopy

Total spheroid area was analysed from light microscopy pictures and measured using the Fiji ImageJ software. For microscopic analysis, spheroids were washed in Dulbecco’s phosphate buffered saline (DPBS, Gibco) after harvesting and subsequently incubated in pre-chilled cell recovery solution (Corning) at 4°C for 20 minutes to remove the Matrigel. The spheroids were washed twice in DPBS, fixed 30 min in 10% formalin (Sigma) and could thereafter be stored in DPBS at room temperature until staining. The exception was spheroids to be stained with Propidium Iodide, which were left unfixed.

Spheroids were stained overnight protected from light in 1:2000 Bodipy-FL (1mg/ml, Invitrogen), 1:500 Dapi (1 mg/ml, Thermo Scientific) and 1:500 Cellmask (5mg/mL, Invitrogen). After the staining, spheroids were washed twice for 5 min in DPBS and then mounted on microscopy slides from iSpacer (SunJin Lab Co) using 90% glycerol as mounting medium. Stained spheroids were analysed by confocal microscopy using a T12 Nikon Eclipse confocal. Equal light settings were used for all images within an experiment and sub-figure. Lipid droplet diameter was measured using the built-in Nikon quantification software, measuring the one largest lipid droplet per adipocyte, in order to have representative data from a mixture of multilocular, paucilocular and unilocular adipocytes. Post-analysis of lipid droplet size was done in GraphPad Prism.

### Measurement of spheroid DNA content

Spheroids were harvested at d30, washed in DPBS, transferred to 200μl new DPBS and centrifuged twice through a Qiagen shredder column at max speed for 2 min to homogenate them. Lysate DNA content was measured using Quant-iT PicoGreen dsDNA Assay Kit (Invitrogen, P7589) according to manufacturer’s instruction.

### Enzyme-linked immunosorbent assay (ELISA) of secreted adipokines

Secretion of the adipokines and cytokines into the cell culture supernatant was measured with ELISA (adiponectin: DLP00, R&D Systems; Leptin: ELH-Leptin-1, Ray Bio; IL-6, ab178013, Abcam; MCP-1, ab179886, Abcam). Leptin samples were diluted 3:4, adiponectin 1:2, MCP-1 1:5, IL-6 1:2. The assay was performed according to manufacturer’s protocol and detected using a plate reader. Post-analysis was done in Excel and grahs in GraphPad Prism.

### Determining lipolysis using the colorimetric glycerol assay

To measure basal and isoprenaline-induced lipid release (lipolysis), each spheroid was washed and subsequently incubated 2 hrs in Krebs-Ringer lipolysis buffer (136 mM NaCl, 1 mM NaH2PO4, 1 mM CaCl2, 4.7 mM KCl, 1mM MgSO4, 2 mM glucose, 25 mM Hepes, 2% fatty acid free BSA, pH 7.4) alone (=basal) or together with 10 μM isoprenaline in a 37°C shaking water bath with lids open. Secreted glycerol levels were measured in the media by mixing 20 μl of sample with 60 μl of Free glycerol reagent (F6428 Sigma) mixed with Amplex UltraRed Reagant at 1:100 and incubating for 15 min at room temperature. The Sigma G7795 standard solution (0.026 mg/mL) was used for standards and absorbance was measured at 540 nm. Post-analysis was done in Excel and graphs in GraphPad Prism.

### Statistical analysis

The data was analysed using one-way ANOVA or linear regression analysis in GraphPad Prism software version 8 (GraphPad Software, San Diego, USA). A p-value of 0.05 was considered significant.

## Results

### Allowing vascular sprouts to guide in vitro adipogenesis

Based on adipocytes differentiating along the microvasculature during development, we reasoned that providing SVF cells with a more natural growth-niche, consisting of vascular endothelial sprouts and extracellular matrix, would help create an adipocyte cell culture model with improved *in vivo*-like cell and tissue morphology. Unsorted human subcutaneous SVF cells were therefore allowed to form floating cell clumps or spheroids by culturing them in ultra-low attachment plates and using endothelial cell media to simultaneously promote endothelial growth^17^ (Fig. 1A). Six days after seeding, spheroids were imbedded in growth factor reduced Matrigel (GFR-Matrigel), shown to be more adipogenic than full Matrigel due to its lack of several anti-angiogenic factors^14^, which resulted in the emergence of clear vascular sprouts within 24 hrs (Suppl. Fig S1A). The GFR-Matrigel-imbedded spheroids were cultured side-by-side with spheroids grown without Matrigel, or traditionally cultured in 2D on imaging slides, neither of these showing any signs of vascular sprouting or tube formation (data not shown). On day 10 (d10) after seeding, half the endothelial media was switched to adipocyte differentiation media, resulting in a rapid increase in spheroid size (Fig. 1B-C). This was especially evident for the Matrigel-imbedded spheroids, where adipogenesis seemed to occur along the vascular sprouts (Fig. 1D-E, compare d10 and d20), becoming much larger than non-imbedded spheroids upon full differentiation on d30 (Suppl. Fig. 1B). The switch to differentiation media induced adipogenesis in all three cultures, as shown by the emergence of multiple Bodipy-stained lipid droplets (d20, Fig. 1E). Remarkably, upon closer microscopic examination, we found on d30 after full differentiation that GFR-Matrigel-imbedding not only resulted in larger spheroids, but also in significantly enlarged adipocyte lipid droplet sizes (Fig. 1E-F). Moreover, the Matrigel-imbedded spheroids more often displayed a unilocular morphology with a single large lipid droplet occupying the central cavity of the cells and the nucleus squashed to the side, whereas in our hands 3D-spheroids cultured without Matrigel had lipid droplets of similar size and morphological distribution as adipocytes cultured in 2D (Fig. 1F). Spheroid size correlated to lipid droplet size throughout all experiments and spheroids cultured with or without Matrigel contained similar amounts of DNA, together suggesting the differences in size between spheroids cultured with or without Matrigel was only due to increased adipocyte size, and not due to differences in preadipocyte proliferation (Fig. 1G-H). For the Matrigel-imbedded spheroids, the average lipid droplet size was somewhat sensitive to the number of seeded SVF cells per spheroid, with too many seeded cells resulting in smaller lipid droplets and 10k cells per spheroid being found to be optimal (Suppl. Fig. 1C). Lastly, our optimized adipocyte spheroid protocol showed very little cell death, evaluated by the absence of both propidium iodide staining and lactate dehydrogenase release to the cell culture media (Suppl. Fig. 1D-E). Adipocyte differentiation was less efficient in the core of the spheroids than in the periphery, potentially due to their large size and poor media diffusion (data not shown). Taken together, by culturing SVF cells in 3D and allowing endogenous, SVF-derived vascular sprouts to form in GFR-Matrigel prior to the induction of adipogenesis, the lipid droplet size and cellular morphology of the differentiated adipocytes were significantly improved as compared to traditional adipocyte culture methods.

**Figure 1.**
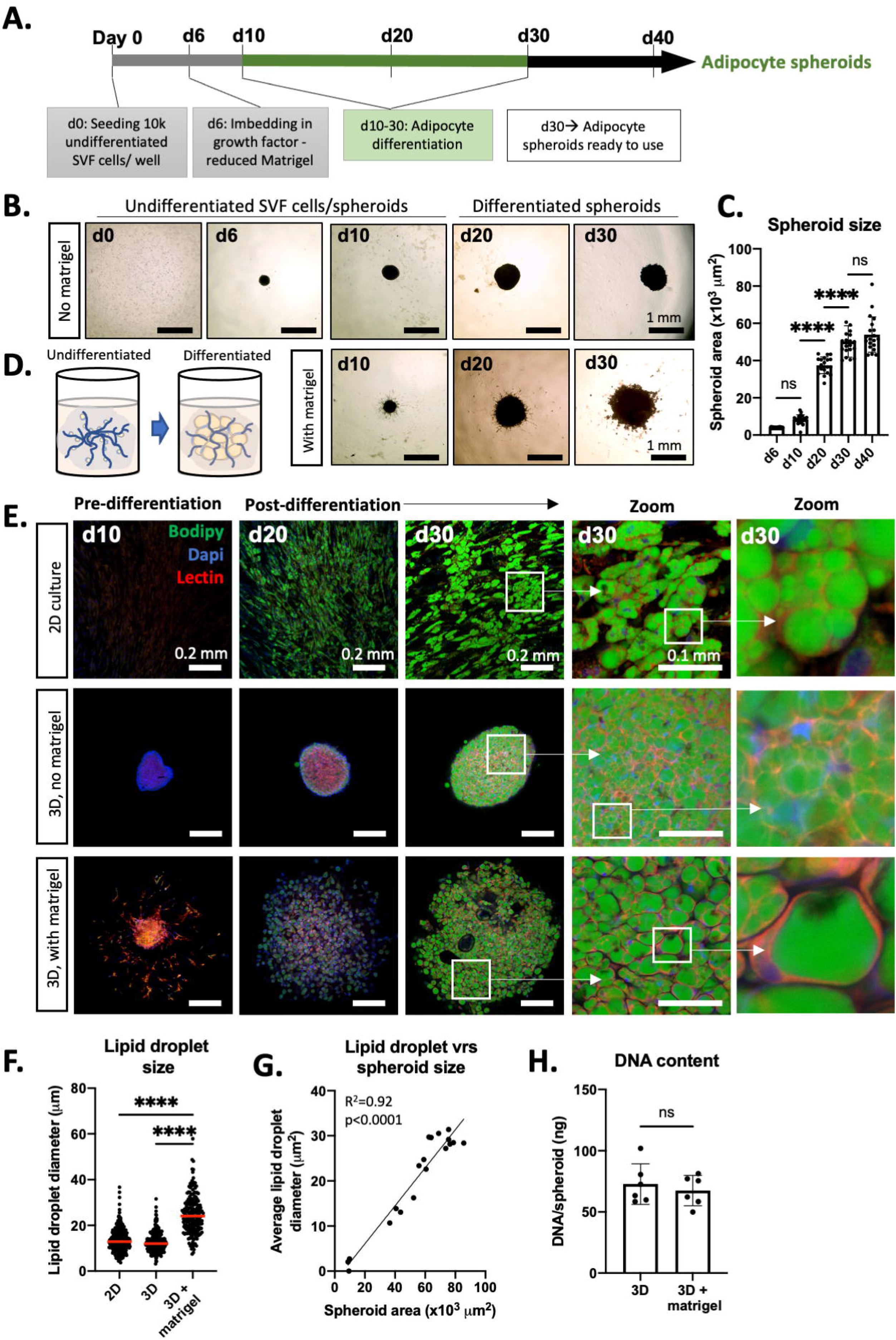
Providing differentiating adipocytes with a vascular growth niche results in larger adipocyte lipid droplets. **A.** Schematic overview of the optimized spheroid culture scheme. **B.** Transmitted light images of spheroids at the indicated days, without or with GFR-Matrigel imbedding. Scale bars 1 mm. **C.** Quantification of the spheroid area at the indicated days, each point is one spheroid, N=20 per time point. **D.** Schematic showing how the vascular sprouts may be contributing as niche for adipocyte differentiation, resulting in larger spheroids. **E.** Representative fluorescent microscopy images of differentiating adipocytes cultured traditionally in 2D (top row), as spheroids without Matrigel imbedding (middle) or with our optimized protocol including Matrigel imbedding (bottom), and stained with Bodipy (green), Dapi (blue) and Lectin (red), showing on the far right the differences in lipid droplet size on d30. **F.** Quantification of lipid droplet sizes for one representative spheroid per culture condition in E. Each point represents one lipid droplet/cell, N= 247 cells in the two upper conditions and N= 203 cells for 3D Matrigel spheroids. **G.** Linear regression analysis of the correlation between average lipid droplet size and spheroid area for N=19 different 3D+Matrigel spheroids at various culture time points, with or without added lipids. **H.** Total DNA content per spheroid, N= 6 spheroids per condition. ****p < 0.0001.

### Characterization of fully differentiated adipocyte spheroids

The morphology and function of the fully differentiated adipocyte spheroids imbedded in GFR-Matrigel were further characterized on d30. Although the average size of the adipocytes and their lipid droplets were significantly smaller as compared to even a lean patient *in vivo*, the morphology and “tissue” architecture was more similar than current 2D models, with adjacent unilocular and paucilocular adipocytes in close contact with each other (Fig. 2A). Compared to previously published 2D and spheroid studies^18,19^ our model showed significantly larger lipid droplet sizes, with mean adipocyte diameter typically reaching 22.9 μm on d30 and 29.1 μm on d40 of culture (Fig. 2B). Upon differentiation (d20 onwards) the spheroids started to secrete high levels of the adipokines adiponectin and leptin into the culture media, confirming full differentiation of the SVF cells into mature adipocytes (Fig. 2C-D). Whereas adiponectin secretion was induced already at d20 and plateaued during the last 10 days of culture, leptin secretion was induced more slowly and correlated with lipid droplet size, as has previously been shown for primary isolated adipocytes^20^ (Fig. 2D-E). The spheroid-associated adipocytes were also responsive to lipolytic cues, responding to isoprenaline stimulation by rapid secretion of glycerol into the surrounding media (Fig. 1F). Importantly, they showed little to no glycerol secretion in the unstimulated state, demonstrating an absence of adipocyte dysfunction despite their larger lipid droplet sizes.

**Figure 2.**
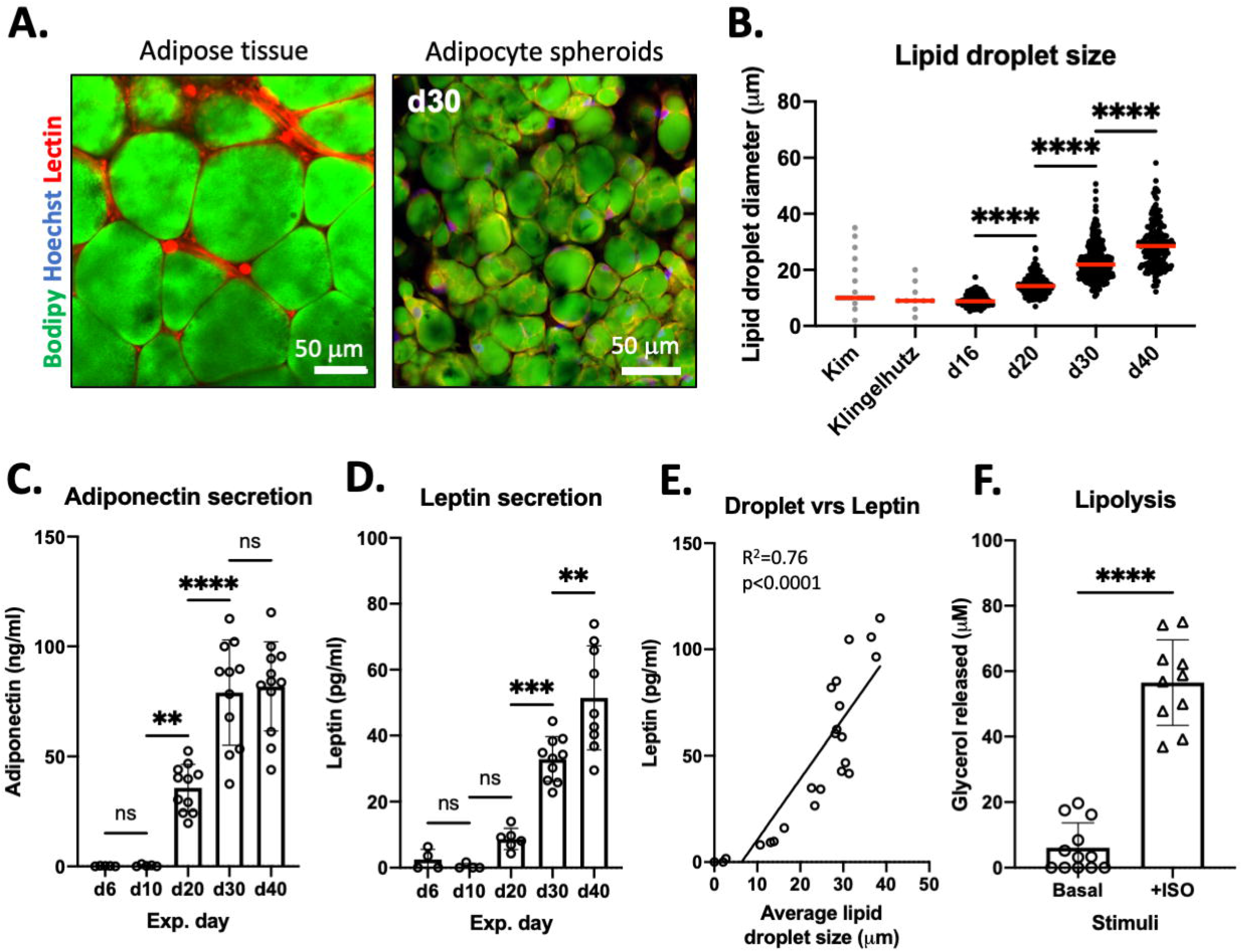
Characterization of fully differentiated adipocyte spheroids. **A.** Comparative images of subcutaneous white adipose tissue from a lean individual, and differentiated subcutaneous adipocytes within our spheroid system, both stained with Bodipy, Lectin and Hoechst. **B.** Quantification of lipid droplet sizes throughout the differentiation process for one representative spheroid per timepoint, each point represents one lipid droplet/cell (N= 107-367 droplets). Data from two references^18,19^ are represented with grey dots to the left. **C-D.** Secretion of adiponectin (c) and leptin (d) to the cell culture media at the indicated days. Each dot represents one spheroid, N= 5 spheroids (d6-10) or 12 spheroids (d20-40). **E.** Linear regression analysis of the correlation between average lipid droplet size and leptin secretion for N=20 spheroids at various culture time points, with or without added lipids. **F**. Basal and isoprenaline (+ISO) stimulated glycerol release during 2 hrs. Each circle (N=12) or triangle (N=10) represents one spheroid. **p < 0.01, ***p < 0.001, ****p < 0.0001.

### Inducing hypertrophy in vitro to mimic weight gain

To date, most mechanistic *in vitro* studies have focused on the effects on adipogenesis, as current *in vitro* models are suboptimal to study mature adipocytes due to the lack of unilocular cells. We therefore set out to test if we could induce lipid droplet enlargement and cellular hypertrophy in our Matrigel-imbedded spheroid system, thus enabling mechanistic studies of mature adipocyte function in response to weight-gain. As a lipid source we chose Intralipid, a soybean oil emulsion used in patient care as a nutritional source of fatty acids or lipid emulsion therapy^21^, because it consists of a more natural mixture of fatty acids and cholesterol than using any single fatty acid alone, and because we reasoned the emulsion would favour lipid uptake into the adipocytes. Spheroids were supplied with 8 mg/dl Intralipid, which is within the normal range of circulating triglyceride concentrations found in plasma. Intralipid-treatment from d10 onwards resulted in significantly increased spheroid sizes at all measured time points (Fig. 3A-C). Subsequent microscopic examination of spheroids stained with Bodipy and lectin showed Intralipid-treatment resulted in on average 25% larger lipid droplets on d30, with average size reaching 30.1 μm (Fig. 3D-E). Interestingly, adding Intralipid at later time points during differentiation, on d14 or d20 instead of at d10, resulted in less of an increase in lipid droplet size, indicating the addition of lipids might enhance differentiation, perhaps through activation of PPARgamma (Suppl. Fig. 1F). In summary, the addition of Intralipid during spheroid differentiation significantly increased adipocyte lipid droplet size, creating a model of inducible hypertrophy *in vitro*.

**Figure 3.**
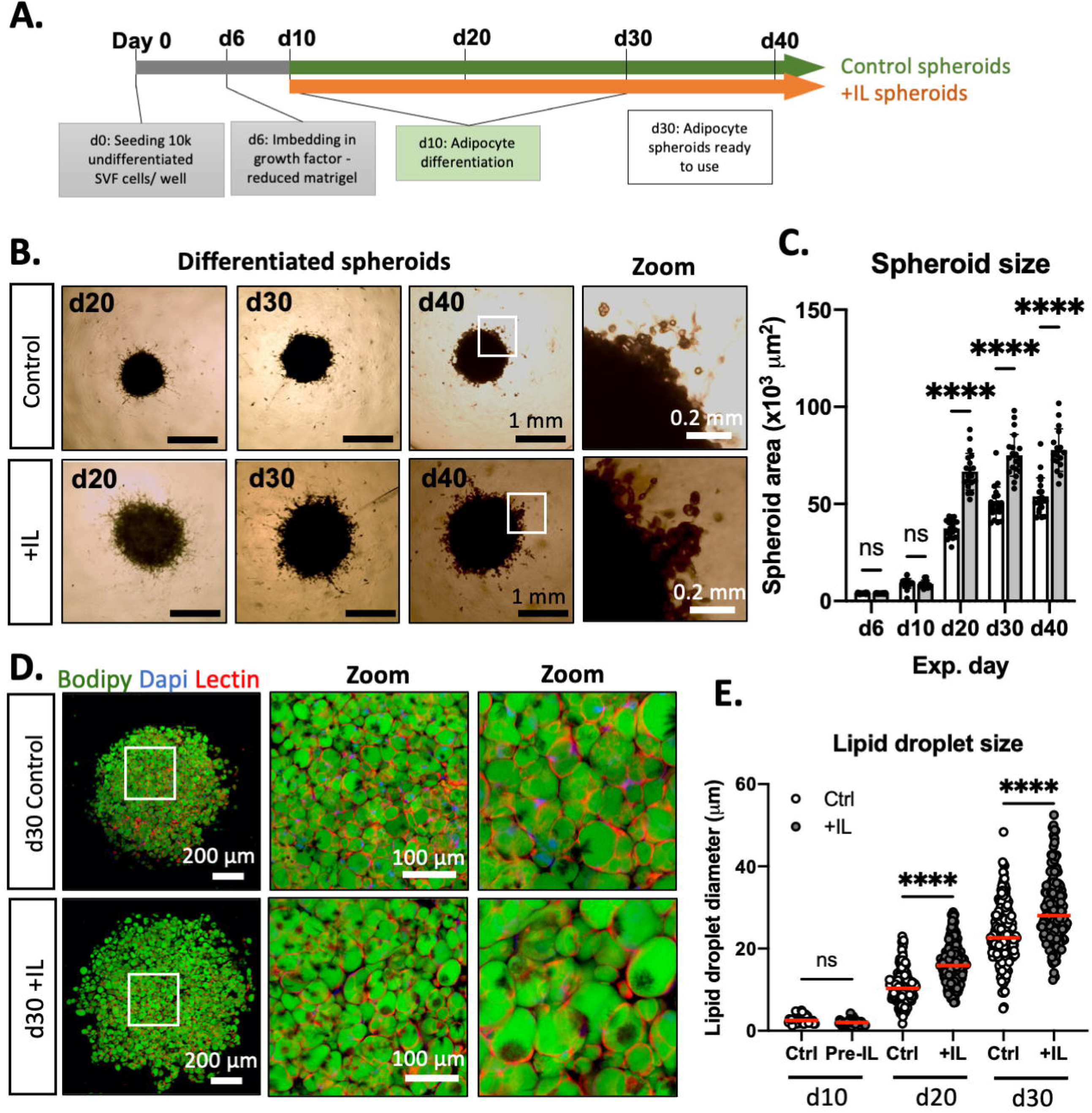
Using Intralipid to induce hypertrophy and mimic weight gain *in vitro*. **A.** Schematic overview of Intralipid (+IL) treatment of spheroids from culture d10 onwards. **B.** Transmitted light images of control and IL-treated spheroids at the indicated days. Zoomed area shows the formation of large round cells on the edges of the spheroids on d40. Scale bar 1 and 0.2 mm, as indicated. **C.** Quantification of the spheroid area for control (white) and IL-treated spheroids (grey) at the indicated days, each point representing spheroid. N=20 spheroids per time point. **D.** Representative fluorescent microscopy images of d30 adipocytes spheroids culture without or with Intralipid (+IL), and stained with Bodipy (green), Dapi (blue) and Lectin (red), showing IL-treated spheroids contain larger lipid droplets. **E.** Quantification of the lipid droplet sizes for one representative control or IL-treated spheroid per day and condition. Each point represents one lipid droplet/cell for N= 42 droplets (d10) or N= 200 droplets (d20-30). ****p < 0.0001.

### Enlarging adipocytes in vitro leads to the development of adipocyte dysfunction

Despite adipocyte dysfunction becoming an established concept in adipocyte biology, it remains unclear what triggers it and why it develops. We asked if merely fattening the adipocytes with Intralipid, resulting in a small but significant increase in adipocyte lipid droplet size (Fig. 3E), would be sufficient to induce characteristics of adipocyte dysfunction, including an altered, proinflammatory adipokine secretion pattern and increased basal lipolysis. Measurement of the secretion of the anti-inflammatory adipokine adiponectin on d30 showed no alterations upon Intralipid-treatment, but in contrast, the secretion of the pro-inflammatory adipokine leptin was found to be significantly induced (Fig. 4A-B). The ratio of adiponectin to leptin secretion was recently proposed to function as a biomarker for adipocyte dysfunction, with lower levels signifying increased dysfunction and development of a more pro-inflammatory obesogenic state^22^. In line with increased adipocyte size associating with adipocyte dysfunction, this ratio was reduced by 67% in Intralipid-treated spheroids as compared to control spheroids (Fig. 4C). Intralipid-treated spheroids moreover secreted higher levels of Interleukin-6 (IL-6) and Macrophage Chemoattractant Protein-1 (MCP-1) to the cell culture media, pro-inflammatory cytokines known to be induced in adipocytes during obesity^5^ (Fig. 4D-E). Lastly, we also measured lipolysis in the fattened spheroids. Whereas basal, non-stimulated lipolysis was found to be increased, the isoprenaline-induced lipolysis was reduced, suggesting the fattened adipocytes had lower responsiveness to external lipolytic cues^6^ (Fig. 4F). Taken together, these tests confirmed that the increase in lipid droplet sizes found in fattened spheroids was accompanied by many of the key characteristics of adipocyte dysfunction, including a lower adiponectin-to-leptin secretion ratio, increased cytokine secretion and dysfunctional lipolysis.

**Figure 4.**
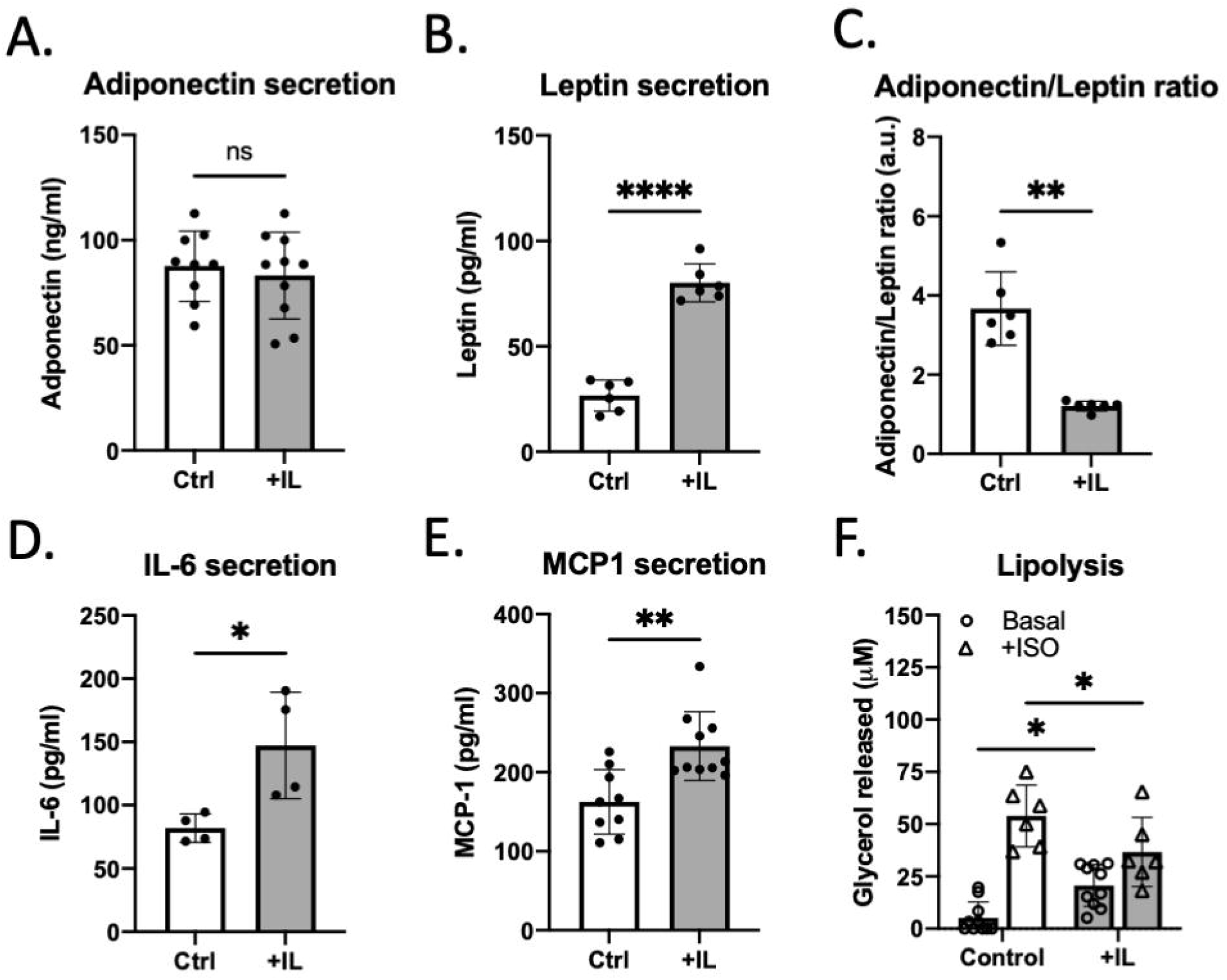
Enlarging adipocytes *in vitro* leads to the development of adipocyte dysfunction. **A-C**. Control and IL-treated spheroid secretion of adiponectin (a) and leptin (b) to the cell culture media on d30, as well as the ratio between the two (c). Each dot represents one spheroid, N= 10 spheroids per condition for adiponectin and 6 spheroids per condition for leptin and the adiponectin/leptin ratio. **D-E.** Control and IL-treated spheroid secretion of IL-6 (d) and MCP-1 (e) to the cell culture media on d30, N= 4 spheroids for IL-6 and 10 spheroids per condition for MCP-1. **F**. Basal (circles) and isoprenaline induced (+ISO, triangles) glycerol release during 2 hrs for control and IL-treated spheroids. Each circle (N=10) or triangle (N=6) represents one spheroid. *p < 0.05, **p < 0.01, ****p < 0.0001.

## Discussion

The rise in obesity prevalence has created an urgent need for new and improved methods to study human adipocytes and the pathogenic effects of weight gain *in vitro*. We present a simple, easily applicable culture protocol that allows for the differentiation and culturing of human adipocytes with a more *in vivo*-like morphology. The protocol moreover uses frozen SVF cells, which today are readily available through commercial vendors, making it applicable to a larger community of laboratories. We also developed a simple protocol to fatten these adipocytes in culture using Intralipid, leading to increased adipocyte lipid droplet size, altered pro-inflammatory signalling and defective lipolysis, all characteristics of adipocyte dysfunction^3,5^. This will enhance further mechanistic studies of the effects of adipocyte dysfunction *in vitro*, and hopefully help advance the obesity research field.

We hypothesize that the spheroid model owes its success to the induction of endogenous vascular sprouts prior to adipocyte differentiation, forming a more optimized growth and differentiation niche for the pre-adipocytes with the SVFs we used (Supl. Fig. S1A). The model builds on previous work by the laboratory of Gou Young Koh, characterizing the importance of the vasculature as niche for postnatal adipocyte differentiation in mice^14^. Undifferentiated pre-adipocytes have also been shown to reside adjacent to micro-vessels within the adipose tissue^23^, making it possible that some preadipocytes migrated along the vascular sprouts into the GFR-Matrigel prior to differentiation, thus explaining the larger spheroid sizes measured for Matrigel-imbedded spheroids as compared to non-imbedded spheroids. It remains to be thoroughly tested if the vascular sprouts are indeed enhancing adipocyte differentiation in our model, and through what signalling pathways this occurs. As such, the spheroid model can be used not only to study mature cells, but also the biology of adipocyte differentiation.

Our model is far from the first adipocyte spheroid. It builds on the knowledge of several other published works showing proof of concept that that culturing adipocyte progenitors in 3D results in larger lipid droplets and tighter cell-cell contacts. Muller et al recently demonstrated that conventional differentiation media kills the SVF endothelial cells, but the use of endothelial medium prior to differentiation can preserve them, resulting in vascularized adipocyte spheroids^17^. They did not, however, focus on characterizing in depth the resulting adipocytes, and it could therefore be interesting to compare their differentiated adipocytes to ours to understand if and how either system could be further improved. Several other studies have also cultured adipocytes as spheroids with or without various types of scaffolds, and although interesting, most reported smaller lipid droplet sizes than those presented here, multilocular morphologies and a sparse “tissue” architecture with poor cellcell contact between adjacent adipocytes^16,18,24,25^. We thus believe the current study offers a significant advancement to previous methods and hope it will contribute to the adipocyte spheroid research community.

Adipocyte enlargement is the primary mechanism for weight gain in humans, and a major focus for obesity research. Despite this, there has to date not existed any optimal *in vitro* method for studying the mechanisms and molecular consequences of adipocyte enlargement in an isolated, controllable cellular environment. Previously, differentiated progenitors cultured in 2D on plastic have been fattened with different fatty acids^19^. Although the saturated fatty acids used in that study were shown to induce lipid droplet enlargement and adipocyte inflammation, the use of a single, saturated fatty acid at high concentrations makes for a synthetic system^26^. By using Intralipid instead, we lose some of the specificity as the exact composition of each batch may vary, but instead supply the adipocytes with a natural mix of lipid species, including high concentrations of both oleic acid and palmitic acids, as well as cholesterol and other lipid species. The emulsified state is moreover believed to contribute to efficient uptake of the lipid mixture to the adipocytes.

Despite the numerous studies showing the advantages of culturing SVFs with either the addition of a matrix or in 3D structures, the majority of studies continue using traditional 2D cultures. We present an easily set-up and relatively streamline method to instead culture SVF cells as Matrigel-imbedded spheroids, resulting in an improved adipocyte morphology and increased lipid droplet sizes. The use of more *in vivo*-like culture models such as this one will result in more physiologically relevant results and biological findings, thus advancing our knowledge of both basic adipose tissue biology as well as the pathological mechanisms acting in response to obesity.

## ACKNOWLEDGEMENTS

We would like to acknowledge the assistance of laboratory technician Anneli Olsson and BSc student Jonathan Ray in the development and culture of adipocyte spheroids. This study was supported by grants from the Swedish Research Council, Jeanssons stiftelse, Åke Wibergs stiftelse, Tore Nilssons stiftelse and Karolinska Institutet.

---

**Supplemental Figure S1.**
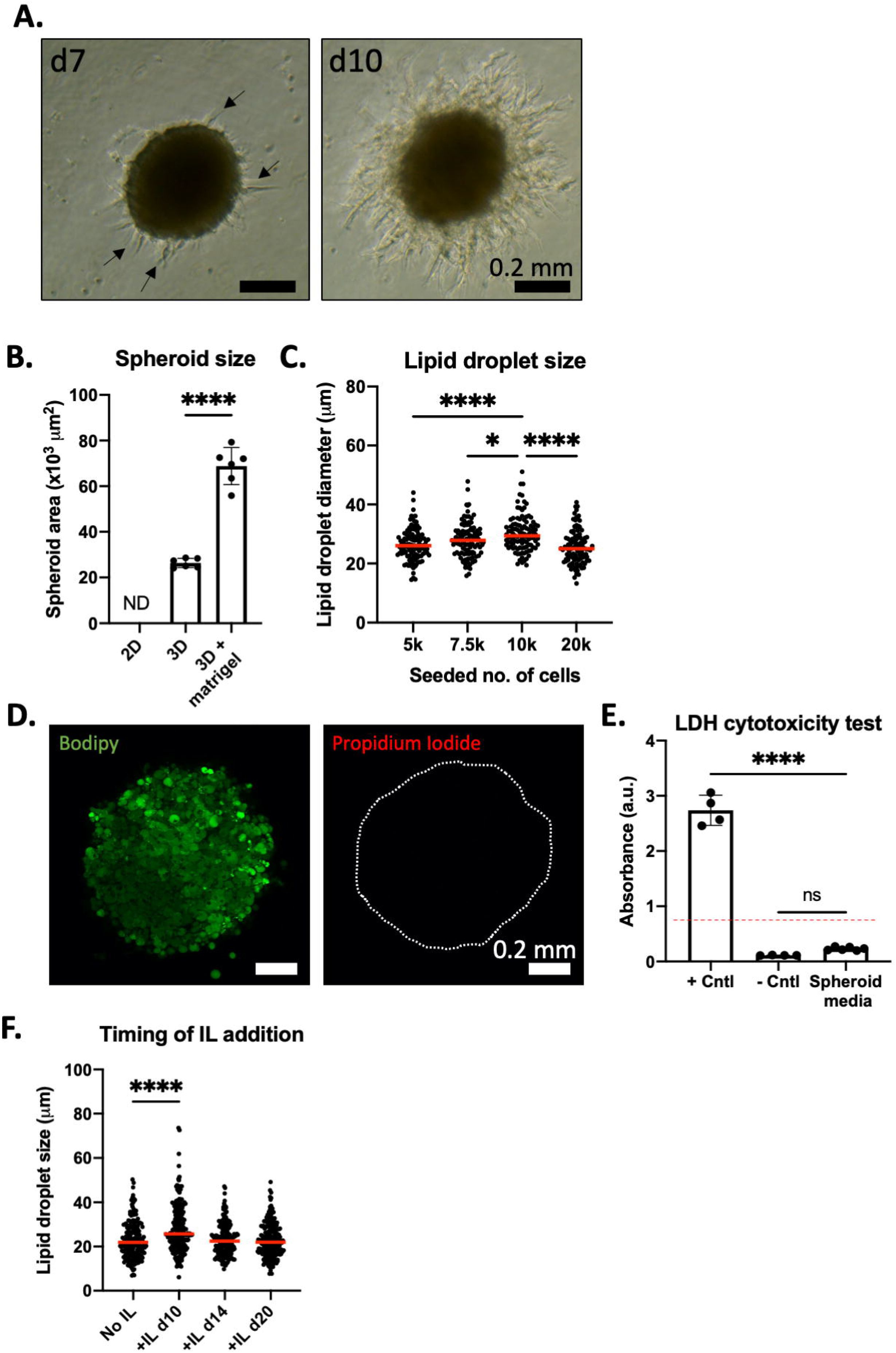
**A.** Enlarged transmitted light images of spheroids at d7 and d10, showing the emergence of vascular sprouts already 24 hrs after GFR-Matrigel imbedding (arrows). Scale bars 0.2 mm. **B.** Quantification of spheroid area for spheroids cultured without or with GFR-Matrigel on d30. Each point is one spheroid, N=6 spheroids per time point. **C.** Quantification of the lipid droplet sizes from two spheroids per condition seeded with the indicated number of SVF-cells per spheroid. Each point represents one lipid droplet size/cell for N= 106 droplets. **D.** Representative fluorescent microscopy images of unfixed, fully differentiated d30 adipocyte spheroids stained with Propidium iodide (red) to detect dead cells, and Bodipy (green). Fixed spheroids were used as positive controls for the staining (data not shown). **E.** LDH-cytotoxicity test detecting release of cytoplasmic content to the spheroid media from dead cells. N= 4 positive or negative control wells and media from N= 6 spheroid wells. Red line denotes limit for cell death detection, above which higher values can be considered positive for cell death. **F.** Quantification of the lipid droplet sizes one representative spheroid per condition, cultured either without Intralipid (no IL) or with IL added at the indicated days of culture (d10, d14 or d20), and lipid droplet sizes measured on d30. Each point represents one lipid droplet size/cell for N= 198 droplets.

## REFERENCES

1. Spalding, K.L., et al. Dynamics of fat cell turnover in humans. Nature 453, 783–787 (2008).

2. Tandon, P., Wafer, R. & Minchin, J.E.N. Adipose morphology and metabolic disease. J Exp Biol 221 (2018).

3. Arner, P., Andersson, D.P., Backdahl, J., Dahlman, I. & Ryden, M. Weight Gain and Impaired Glucose Metabolism in Women Are Predicted by Inefficient Subcutaneous Fat Cell Lipolysis. Cell Metab 28, 45–54 e43 (2018).

4. Longo, M., et al. Adipose Tissue Dysfunction as Determinant of Obesity-Associated Metabolic Complications. Int J Mol Sci 20 (2019).

5. Hammarstedt, A., Gogg, S., Hedjazifar, S., Nerstedt, A. & Smith, U. Impaired Adipogenesis and Dysfunctional Adipose Tissue in Human Hypertrophic Obesity. Physiol Rev 98, 1911–1941 (2018).

6. Ryden, M., Andersson, D.P., Bernard, S., Spalding, K. & Arner, P. Adipocyte triglyceride turnover and lipolysis in lean and overweight subjects. J Lipid Res 54, 2909–2913 (2013).

7. Acosta, J.R., et al. Increased fat cell size: a major phenotype of subcutaneous white adipose tissue in non-obese individuals with type 2 diabetes. Diabetologia 59, 560–570 (2016).

8. Kusminski, C.M., Bickel, P.E. & Scherer, P.E. Targeting adipose tissue in the treatment of obesity-associated diabetes. Nat Rev Drug Discov 15, 639–660 (2016).

9. Hagberg, C.E., et al. Flow Cytometry of Mouse and Human Adipocytes for the Analysis of Browning and Cellular Heterogeneity. Cell Rep 24, 2746–2756 e2745 (2018).

10. Verboven, K., et al. Abdominal subcutaneous and visceral adipocyte size, lipolysis and inflammation relate to insulin resistance in male obese humans. Sci Rep 8, 4677 (2018).

11. Harms, M.J., et al. Mature Human White Adipocytes Cultured under Membranes Maintain Identity, Function, and Can Transdifferentiate into Brown-like Adipocytes. Cell Rep 27, 213–225 e215 (2019).

12. Skurk, T. & Hauner, H. Primary culture of human adipocyte precursor cells: expansion and differentiation. Methods Mol Biol 806, 215–226 (2012).

13. Yu, J. & Li, P. The size matters: regulation of lipid storage by lipid droplet dynamics. Sci China Life Sci 60, 46–56 (2017).

14. Han, J., et al. The spatiotemporal development of adipose tissue. Development 138, 5027–5037(2011).

15. Cho, C.H., et al. Angiogenic role of LYVE-1-positive macrophages in adipose tissue. Circ Res 100, e47–57 (2007).

16. Louis, F., Kitano, S., Mano, J.F. & Matsusaki, M. 3D collagen microfibers stimulate the functionality of preadipocytes and maintain the phenotype of mature adipocytes for long term cultures. Acta Biomater 84, 194–207 (2019).

17. Muller, S., et al. Human adipose stromal-vascular fraction self-organizes to form vascularized adipose tissue in 3D cultures. Sci Rep 9, 7250 (2019).

18. Klingelhutz, A.J., et al. Scaffold-free generation of uniform adipose spheroids for metabolism research and drug discovery. Sci Rep 8, 523 (2018).

19. Kim, J.I., et al. Lipid-overloaded enlarged adipocytes provoke insulin resistance independent of inflammation. Mol Cell Biol 35, 1686–1699 (2015).

20. Jernas, M., et al. Separation of human adipocytes by size: hypertrophic fat cells display distinct gene expression. FASEB J 20, 1540–1542 (2006).

21. Muller, S.H., Diaz, J.H. & Kaye, A.D. Clinical applications of intravenous lipid emulsion therapy. J Anesth 29, 920–926 (2015).

22. Fruhbeck, G., Catalan, V., Rodriguez, A. & Gomez-Ambrosi, J. Adiponectin-leptin ratio: A promising index to estimate adipose tissue dysfunction. Relation with obesity-associated cardiometabolic risk. Adipocyte 7, 57–62 (2018).

23. Tang, W., et al. White fat progenitor cells reside in the adipose vasculature. Science 322, 583–586 (2008).

24. Abbott, R.D., et al. The Use of Silk as a Scaffold for Mature, Sustainable Unilocular Adipose 3D Tissue Engineered Systems. Adv Healthc Mater 5, 1667–1677 (2016).

25. Graham, A.D., et al. The development of a high throughput drug-responsive model of white adipose tissue comprising adipogenic 3T3-L1 cells in a 3D matrix. Biofabrication 12, 015018 (2019).

26. Al-Sulaiti, H., et al. Triglyceride profiling in adipose tissues from obese insulin sensitive, insulin resistant and type 2 diabetes mellitus individuals. J Transl Med 16, 175 (2018).

